# TurboID-based proximity labeling enables *in vivo* mapping of *Plasmodiophora brassicae* secretome in Arabidopsis

**DOI:** 10.1101/2025.11.18.684874

**Authors:** Kris Kalinger, Emily Gallipeau-Burns, Md. Musharaf Hossain, Maryam Nourimand, Elzbieta Mietkiewska, Mohana Talasila, R. Glen Uhrig, Christopher D. Todd, Allyson M. MacLean

## Abstract

- *Plasmodiophora brassicae*, the causal agent of clubroot disease, is an obligate biotrophic protist belonging to the poorly characterized *Rhizaria*. Its intracellular lifestyle and resistance to genetic manipulation have hindered functional analysis of its effector repertoire, leaving mechanisms underlying disease development unresolved. Here, we sought to experimentally define the *P. brassicae* secretome within infected plant cells and identify effectors targeted to specific host subcellular compartments.
- A proximity labeling approach based on the TurboID biotin ligase was used to capture pathogen-derived proteins within the nucleus, cytosol, endoplasmic reticulum, and plasma membrane of infected Arabidopsis roots during primary and secondary stages of clubroot disease.
- This strategy yielded the first *in planta* experimental view of the *P. brassicae* secretome, identifying both established and previously uncharacterized effectors. The resulting dataset provides a valuable resource and methodological framework for dissecting effector function in this and other intracellular plant pathogens.
- This study expands our understanding of *Rhizarian* pathogenicity and provides a methodological template for identifying the secretomes of other obligatory intracellular plant pathogens.

## Introduction

*Plasmodiophora brassicae* is an obligate biotrophic pathogen and the causal agent of clubroot disease in members of the *Brassicaceae* family (Adhikary *et al*., 2025). Its host range includes the model plant *Arabidopsis thaliana* (Koch *et al*., 1991) as well as economically important crops such as *Brassica napus* L. (rapeseed, canola) and *Brassica oleracea* (cabbage) (Dixon, 2009). In susceptible hosts, disease progression is marked by the development of increasingly thickened and deformed roots, culminating in the formation of characteristic club-shaped galls. These alterations in root architecture have been attributed to *P. brassicae*–induced reprogramming of host developmental pathways and physiology, which collectively establish a nutrient sink to support proliferation of the protist (Malinowski *et al*., 2019; Vañó *et al*., 2023; Blicharz *et al*., 2024), and the production of resting spores that comprise the next generation. As the disease advances, vascular development becomes progressively dysregulated (Malinowski *et al*., 2012), impairing the plant’s capacity to absorb water and nutrients from the soil and leading to visible symptoms in aboveground tissues such as stunting, wilting, and yield loss. With time, the infected root galls undergo senescence, eventually releasing resting spores into the soil to initiate a new round of infection. Numerous major resistance (CR) genes and quantitative trait loci (QTLs) have been identified in *Brassica* species (e.g., *B. rapa*, *B. oleracea*, *B. napus*) that confer resistance against specific pathotypes of *Plasmodiophora brassicae* (Manzanares-Dauleux *et al*., 2000; Suwabe *et al*., 2003; Hirai *et al*., 2004; Matsumoto *et al*., 2012; Hatakeyama *et al*., 2013; Chu *et al*., 2014; Hasan & Rahman, 2016; Farid *et al*., 2020). However, there is growing evidence that resistance in agronomically important crops such as canola is breaking down (Strelkov *et al*., 2016; Mcdonald *et al*., 2021). This underscores the need to improve our understanding of the molecular mechanisms by which *P. brassicae* manipulates its hosts, particularly through the action of secreted effector proteins, to guide the development of more durable strategies for disease management.

A soil-borne protist, *P. brassicae* initiates a primary stage of infection by entering epidermal and root hair cells, with a subsequent secondary stage of infection within the root cortical cells, where the protist induces extensive cell hypertrophy and proliferation (Liu *et al*., 2020). These physiological changes are central to the formation of root galls and are most likely driven, at least in part, by pathogen-secreted effector proteins that manipulate host cellular processes to facilitate infection and nutrient acquisition (Pérez-López *et al*., 2018). Despite its agronomic significance, molecular studies of *P. brassicae* have been limited by an obligate lifestyle that precludes routine culture and genetic manipulation (Dixon, 2014). As a result, and despite significant progress in this area over the last decade (Chen *et al*., 2019; Hossain *et al*., 2021; Pérez-López *et al*., 2021; Ando *et al*., 2024; Yang *et al*., 2024), it is likely that the identity, localization, and mode of action of many *P. brassicae* effectors remain unknown. Conventional effector discovery pipelines which typically integrate whole-genome or transcriptome sequencing with bioinformatic predictions of N-terminal secretion signals, provide a foundational approach for identifying candidate effectors (Zhan *et al*., 2022). However, these methods are constrained by underlying assumptions regarding canonical secretion pathways and well-characterized modes of host cell entry derived from other pathosystems that remain largely untested in parasitic protists such as *P. brassicae*. Unbiased, experimentally driven approaches are therefore needed to define the full effector repertoire deployed during clubroot infection.

Proximity labeling methods combined with mass spectrometry (PL-MS), including the use of engineered biotin ligases such as TurboID (Branon *et al*., 2018), offer a powerful, high-resolution approach to study protein interactions *in planta* (Xu *et al*., 2023). TurboID enables *in vivo* biotinylation of proteins within a ∼10 nm radius of the enzyme (Kim *et al*., 2014), allowing for the identification of transient or low-abundance interactions under native physiological conditions (Mair *et al*., 2019). This approach has been successfully employed to characterize host-pathogen interfaces in bacterial and fungal systems (Shi *et al*., 2023) but has yet to be applied within the context of biotrophic protist pathogens, such as *P. brassicae*. In this study, we apply TurboID-mediated proximity labeling to identify *P. brassicae* proteins that are delivered into host cells during infection. By expressing TurboID in specific subcellular compartments of infected *Arabidopsis thaliana* roots and capturing biotinylated proteins during critical stages of primary infection and root gall development, we aim to define a set of proteins secreted by this pathogen during host colonization. This approach provides a platform for the unbiased discovery of proteinaceous effectors, further advancing our understanding of *P. brassicae* pathogenesis and host manipulation strategies.

## Materials and Methods

### Plant materials and growth conditions

All seeds were surface-sterilized prior to sowing by incubation for 1m in 70% v/v ethanol, followed by 7m in 50% v/v household bleach (3% sodium hypochlorite) with 0.01% Tween-20. Seeds were rinsed 5 times with sterile distilled water following surface sterilization.

*Arabidopsis* seeds were stratified in the dark at 4°C for three days prior to sowing on autoclaved Pro-Mix BX loose soil medium. Plants were then grown in a Conviron ATC40 growth chamber under long-day (16h light / 8h dark) conditions at 22°C / 18°C day/night temperatures, 150 µmol / m^2^ light intensity, and 60% relative humidity. To facilitate harvesting root material, *Arabidopsis* plants used for inoculation with *P. brassicae* and biotin treatments were transplanted 10 days after germination to 50-cell trays containing a 1:1 mixture of sterilized sand and Turface® calcinated clay, one plant per cell. Plants were kept well-watered and fertilized every three weeks with 10mL 1/4-strength 20-20-20 Miracle-Gro® all purpose plant food.

Wild type *N. benthamiana* seeds were sown on Sunshine Mix #3 soil (Sun Gro Horticulture Inc., Vancouver, BC, Canada) and kept at 4 °C for two days for cold stratification. Seeds germinated at 25 °C under 16 hours of light and 8 hours of dark, with a light intensity of 160 µmol photons m⁻² sec⁻¹ in a growth chamber (Conviron E8, CMP6050 control system). After ten days post germination, each seedling was transplanted into individual pots for growth. Leaves from five-week-old *N. benthamiana* plants were then used for transient expression of the generated constructs in subcellular co-localization studies.

### Plasmid construction

All constructs were assembled in the binary vector pCAMBIA2301 using Gibson Assembly or traditional restriction digest and ligation. As a first step, the native *35S:GUS* cassette in pCAMBIA2301 was excised with XbaI and PmlI to generate a promoterless backbone.

#### EXP7p:GFP-NES

A 774 bp *eGFP* fragment was amplified from *eGFP*/DONOR2-3 using primers EM-F1/EM-R1 to add an N-terminal FLAG tag and C-terminal NES with two stop codons, then re-amplified with EM-F4/EM-R4 for Gibson overlaps.

The *EXP7* promoter (1.4 kb) was amplified from N2106306 (NASC) with EM-F3/EM-R3 and assembled with *FLAG-eGFP-NES* into the XbaI/PmlI sites of pCAMBIA2301 to yield *EXP7p:GFP-NES*. To create ***EXP7p:TurboID-NES***, a 1.0 kb *TurboID-NES* fragment was amplified from pENTR_L1-YFP-Turbo-NES-L2 (Addgene #127349) using EM-F2/EM-R2, re-amplified with EM-F4/EM-R4, and assembled with the same *EXP7* promoter.

For all other plasmids, promoter-specific backbones were created by inserting either the 1.4 kb *EXP7* or 1.5 kb *PEP* promoter into XbaI/PmlI-cut pCAMBIA2301 following PCR amplification using primers F13/R13 and F14/R14, respectively, to generate EM7 (EXP7-pCAMBIA2301) and EM8 (PEP-pCAMBIA2301). These served as universal cloning bases for subsequent assemblies into the PmlI site.

### Cytosolic and ER-Localized Constructs

*PEPp:GFP-NES* and *PEPp:TurboID-NES* were obtained by inserting FLAG-tagged *eGFP* or *TurboID* fragments (amplified by primers sets EM-F8/EM-R4) into PmlI-linearized EM8. ER-retained constructs were generated similarly, with PCR products cloned into linearized EM7 or EM8 via PmlI sites: *EXP7p:GFP-KDEL*, following amplification with EM-F4/EM-R8, *EXP7p:TurboID-KDEL*, following amplification with EM-F4/EM-R9a, and *PEPp:GFP-KDEL* or *PEPp:TurboID-KDEL*, following amplification using EM-F11/EM-R10 to introduce promoter overlaps.

### CERK1-Based Membrane-Localized Constructs

The CERK1 CDS was amplified from *A. thaliana* Col-0 cDNA with EM-F46/EM-R46 and cloned into pCR-Blunt II-TOPO, yielding EM29. Derived fusions were subsequently assembled in EM7 or EM8 backbones using Gibson Assembly.

To generate *EXP7p:TM_CERK1_–TurboID*, a CERK1 fragment containing the Kozak sequence was amplified from EM29 with EM-F49 and EM-R49, while the TurboID fragment was amplified from *EXP7p:TurboID-NES* using EM-F50 and EM-R50, followed by reamplification with EM-F50 and EM-R51 to introduce a C-terminal FLAG epitope, dual stop codons, and overlap with the NOS terminator. The *TM_CERK1_–TurboID* fragment was cloned into the PmlI site of EXP7-Cambia2301 (EM7 plasmid) using the Gibson assembly approach.

To create *EXP7p:GFP-TM_CERK1_–TurboID*, a four-fragment Gibson assembly was performed using *CERK1*, *eGFP*, *CERK1 transmembrane linker*, and *TurboID* fragments amplified with EM-F49/EM-R52, EM-F53/EM-R53, EM-F54/EM-R54, and EM-F55/EM-R55, respectively, cloned into the PmlI site of EM7.

Similarly, *EXP7p:TM_CERK1_–GFP* was assembled using *CERK1* and *eGFP* fragments amplified with EM-F49/EM-R56, EM-F57/EM-R57, and reamplified with EM-F57/EM-R51 to add vector overlaps, and cloned into PmlI site of EM7.

The *PEP*-driven counterparts were constructed analogously: PCR amplification was performed for *PEPp:TM_CERK1_–TurboID* using EM-F58/EM-R51, *PEPp: GFP–TM_CERK1_–TurboID* using EM-F58/EM-R55, and *PEPp:TM_CERK1_–GFP* using EM-F58/EM-R51. Products were then cloned into PmlI site of EM8.

All plasmids were verified by DNA sequencing and primer sequences are listed in Table S1.

### Generation of transgenic plants (text from Ela)

Transformation of Arabidopsis (ecotype Columbia) with the developed constructs was performed using a floral dip method as described by (Clough & Bent, 1998). Arabidopsis T1 seeds transformed with the described vectors were screened on MS media (Murashige & Skoog, 1962) supplemented with Kanamycin (50 mg/L). Kanamycin resistant, T1 plants were grown in soil to develop T2 seed generation. All T2 transgenic seed lines were subjected to segregation analysis to identify lines with transgene insertion at one locus (segregation 3:1 on selection media). Select T2 seed lines with transgene insertion at one locus were grown in soil to develop T3 seed generation which were screened to select homozygous lines (Mietkiewska *et al*., 2007).

### Maintenance of P. brassicae and inoculation of Arabidopsis

*P. brassicae* was propagated using the canola (*Brassica napus*) cultivar ‘Westar’ as a host. Plants were cultivated in a greenhouse with an ambient daytime temperature range of temperature ranged 22-26°C and nighttime temperature range of 18-23°C. The relative humidity was maintained at 40-70%. Surface-sterilized canola seeds were sown in a starter tray of dampened, sterilized vermiculite, then transferred 7 days after germination to 50-cell trays containing Pro-Mix BX loose soil medium, one plant per cell. 1 week later, plants were inoculated with 2 × 10^8^ spores / mL of *P. brassicae* resting spore suspension, prepared from frozen stocks of canola root galls according to (Salih *et al*., 2024). Three days post-inoculation, plants were moved to individual 5-inch pots containing Pro-Mix BX loose soil medium. Plants were fertilized once every 2 weeks by bottom watering with full-strength 20-20-20 Miracle-Gro® all-purpose plant food. Infected root galls were harvested from inoculated plants 4-5 weeks post-inoculation, when large galls had formed. Galls were stored at 20°C until used for resting spore isolation, for up to 3 months. Ten-day-old *Arabidopsis* plants transplanted to a 1:1 sand:Turface® mixture were allowed to acclimate for 3 days prior to inoculation with *P. brassicae* resting spores. Resting spores were harvested from canola root galls according to (Salih *et al*., 2024) and the spore suspension was diluted with sterile water to 1 × 10^6^ spores / mL. The fresh suspension was used immediately for plant inoculation. Plants were watered thoroughly, then inoculated by applying 1mL of *P. brassicae* spore suspension around the base of each plant.

### Subcellular localization assays in N. benthamiana

To determine the subcellular localization of constructs driven by the 35S promoter, *A. tumefaciens* GV3101 containing the desired construct was transiently co-expressed with organelle-specific markers in leaves of five-week-old *N. benthamiana* plants. The NLS-mCherry served as a nuclear localization marker to validate NLS/NES constructs, while mCherry-HDEL (ABRC: CD3-959) marked the ER. The final OD600 was adjusted to 0.5 for each construct. For co-localization studies, a mixture of *A. tumefaciens* GV3101 containing the organelle marker and the construct, at equal concentrations, was infiltrated into the abaxial side of *N. benthamiana* leaves. The subcellular localization of the fluorescent-tagged protein was examined two days after agroinfiltration using a confocal laser scanning microscope (LSM880 with Airyscan; ZEISS). GFP fluorescence was excited with an Argon laser at 488 nm, and emissions were captured between 500 and 530 nm, whereas mCherry tags were excited at 561 nm with a Helium-Neon laser, and emissions were collected between 600 and 630 nm. To prevent overexpression artifacts, cells with relatively low fluorescence signals were imaged after transient expression and later analyzed with ImageJ (https://imagej.net/Fiji).

### Subcellular localization assays in Arabidopsis

Surface-sterilized Arabidopsis seeds were sown on square plates containing solid 1/2 Murashige and Skoog basal medium and stratified in the dark at 4°C for 3 days. The plates were then moved to a Panasonic MLR-352H-PA growth chamber under long-day (16h light / 8h dark) conditions at 22°C / 18°C day/night temperatures, 150 µmol / m^2^ light intensity, and 60% relative humidity. Roots were cut into approximately 1 cm lengths and placed on slides (Fisher Scientific, 25 x 75 x 1.0 mm) with a drop of water, then a cover slip (Globe Scientific Inc. 22 x 22 mm, #1. 5) (0.17 μm) was placed on top. Samples were observed and fluorescence was imaged using a Nikon A1RsiMP laser scanning confocal microscope with a 10x dry objective lens (Plan APO λ 0.45). eGFP (488 nm) was excited using an Aragon laser, and emitted fluorescence was captured from 500-550 nm for eGFP. Brightfield images were captured simultaneously with fluorescence using a Nikon A1 Plus camera. Z-stack projections were generated using the Galvano scanner, with each step in the stack being 2.1 μm. Images were processed using FIJI (Schindelin *et al*., 2012).

To obtain images of root hair infection by motile *P. brassicae* zoospores, 1mL of a freshly prepared resting spore suspension, diluted to 1 × 10^4^ spores / mL, was spread across the surface of MS plates. Roots were harvested for imaging 24h and 48h post-inoculation.

### Biotin activity assays and affinity purification of biotinylated proteins

For plants expressing PEP_p_ constructs, plants were grown for 4 weeks post-inoculation with *P. brassicae* resting spores to allow for the formation of root galls prior to biotin treatment, while plants expressing EXP7_p_ constructs were treated with biotin 48h post-inoculation. Biotin treatment consisted of watering each plant with 20mL of a 200µM biotin solution, or with 0.005% DMSO in water as a control. Excess solution was drained from the bottom of the trays, after which the biotin labelling reaction was left to proceed for 10h. Plants were then uprooted and placed in ice water to halt the labelling reaction, and either whole root systems (plants expressing EXP7_p_ constructs) or excised gall tissue (plants expressing PEP_p_ constructs) were harvested and flash-frozen in liquid nitrogen. Pooled root material from 25 plants constituted a single biological replicate (n = 25).

Protein extraction and streptavidin affinity purification was then conducted according to Mair *et al*. (2019) with some modifications. Frozen, infected root material was weighed, homogenized in liquid nitrogen, resuspended in 2 volumes of ice-cold extraction buffer (50mM Tris pH 7.5, 150mM NaCl, 0.1% SDS, 1% Triton X-100, 0.5% sodium deoxycholate, 1mM EDTA pH 7.5, 1mM DTT, 1mM PMSF, 1 Roche Complete™ Protease Inhibitor Cocktail tablet / 100mL, 0.1 mg / mL lysozyme from chicken egg white, 5U DNAse I / 1mL) and incubated at 4°C on a shaker platform for 15 minutes. Samples were then sonicated on ice for three 20-second cycles, with 90s between each cycle. and centrifuged for 5 minutes at 5,000 x *g* and 4°C to pellet cell debris. The supernatant, representing the crude protein extract, was passed through one or more disposable PD-10 desalting columns (max vol. 2.5mL extract / column) to remove excess biotin. Columns were equilibrated according to manufacturers’ instructions with 25mL ice-cold equilibration buffer (50mM Tris pH 7.5, 150mM NaCl, 0.1% SDS, 1% Triton X-100, 0.5% sodium deoxycholate, 1mM EDTA pH 7.5, 1mM DTT) prior to use, and 3.5mL of extraction buffer was used to elute 2.5mL of protein extract from each column. A small portion of eluate was retained for immunoblotting analysis to confirm biotinylation activity, and for determination of total sample protein concentration using Pierce™ Bradford Plus Protein Assay Reagent (Thermo Fisher), according to manufacturers’ instructions. Samples were diluted 1:5 to prevent detergents in the sample extraction buffer from interfering with the Bradford assay. For every 4g total protein, 50µL of High-Capacity Magne ® streptavidin-conjugated bead slurry (Promega) was added to the sample extract. The beads were washed 3 times with 1mL ice-cold extraction buffer prior to use. After the addition of streptavidin beads, PMSF and dissolved Complete ™ Protease Inhibitor Cocktail were added to each sample to reach a 1X final concentration of each, and samples were incubated at 4° C for 16h on a shaker platform. The next day, the beads were separated from the supernatant using a magnetic rack, and a portion of the supernatant was retained for immunoblotting to confirm success of the affinity purification procedure. The beads were washed with several solutions according to the protocols of Mair *et al*. (2019) and Branon *et al*. (2018) to remove any weakly bound proteins. The beads were then resuspended in 2 volumes of equilibration buffer. Approximately 10% of each bead sample was boiled in 4X Laemmli buffer containing 20mM DTT and 2mM biotin, at 95°C for 5 minutes, and used for immunoblotting analysis. The remaining beads were stored at −80°C for MS sample preparation.

### SDS-PAGE and Immunoblots

To screen the labelling efficiency of TurboID-expressing transgenic plant lines, frozen, homogenized, biotin-treated root material was prepared for immunoblotting by boiling in 1 volume of 1X Laemmli buffer (60 mM Tris pH 6.8, 2% SDS, 10% glycerol, 20mM DTT, 0.025% bromphenol blue) for 5min at 95°C. Crude protein extracts and unbound protein samples from affinity purification were prepared for immunoblotting by boiling in 1 volume of 2X Laemmli buffer. Streptavidin bead-bound proteins were dislodged from the beads as described previously. Proteins were separated by SDS-PAGE and transferred onto 0.45µm nitrocellulose membrane for 25min at 1.3A using a Trans-Blot Turbo semi-dry transfer apparatus (BioRad). Immunoblots were blocked in 3% w/v bovine serum albumin in TBS-T (20mM Tris pH 7.6, 150mM NaCl, 0.1% Tween-20) since commonly used milk blocking buffer contains many biotinylated proteins that interfere with streptavidin immunoblotting. Streptavidin-HRP (S911, Thermo Fisher Scientific) was used to probe blots for biotinylated proteins, and mouse monoclonal anti-GFP HRP-conjugated primary antibody (GF28R HRP, Thermo Fisher Scientific) was used to detect the presence of GFP in control samples. Blots were probed with overnight on a shaker platform at 4°C, and incubated with Pierce™ ECL Western Blotting Substrate (Thermo Fisher Scientific) according to the manufacturer’s instructions. Blots were imaged on a BioRad ChemiDoc MP imaging system.

### Sample preparation, Liquid Chromatography Mass Spectrometry analysis and data processing

#### Sample Preparation

Protein bound streptavidin beads were washed sequentially twice with 1x wash buffer 1 (50 mM HEPES pH 7.5, 150 mM NaCl, 0.5% (v/v) Triton X-100, 0.5% (v/v) NP-40), 1x wash buffer 2 (1 M KCl), 1x buffer 3 (100 mM Na_2_CO_3_), 1x wash buffer 4 (2M urea in 50mM Tris-HCl pH 8.0), 3x buffer 5 (50 mM triethylammonium bicarbonate (TEAB) in HPLC water). The washed, sample-bound beads were then digested with 1 µg Trypsin (Promega; V5113) in 100uL 50 mM TEAB in HPLC water at 37°C overnight (16 h) using a modified protocol on Kingfisher Apex (ThermoFisher Scientific) from (Leutert *et al*., 2019). Peptides were then sequentially reduced (10 mM Dithiothreitol) at 95°C for 5 min, cooled to room temperature, and alkylated (30 mM Iodoacetamide) for 30 minutes in the dark, then acidified with 25% (v/v) Trifluoroacetic acid to a final concentration of 0.5% (v/v) and pH < 2. Acidified peptides were then desalted using OMIX C18 tips (Agilent; A5700310K) mounted on Opentrons OT-2 pipetting robot. The desalted phosphopetides were dried in a SpeedVac (Thermo) and stored at −80℃ until re-suspension in 3% (v/v) ACN and 0.1% (v/v) Formic Acid (FA) for LC-MS analysis.

#### Mass Spectrometry Analysis

The total proteome peptide pool was analyzed using a 15 cm Easy-Spray PepMap C18 Column (Thermo Scientific; ES904) heated to 50°C with a flow rate of 300nL/min. Mass spectrometry data was acquired using a data dependent acquisition, where the peptides were eluted at 300 nL/min over a 120-minute gradient from 4 - 41% solvent B (0.1% (v/v) formic acid; FA in 80% Acetonitrile; ACN) as previously described (Mehta *et al*., 2022).

#### Data Processing

Raw mass spectrometry data was analysed Using MaxQuant ver. 2.5.1.0 with default settings, including: Carbidomethyl (C) as a fixed modification, methionine oxidation (M) as a variable modification, a maximum of 2 missed cleavages and peptide spectrum match (PSM), peptide, and protein group FDR < 0.01. Data was searched using either the ARAPORT 11 *Arabidopsis thaliana* proteome or the *Plasmodiophora brassicae* proteome (GCA_001049375), respectively.

## Results

### Designing a proximity labeling platform to profile the P. brassicae secretome

As a means of enabling detection and subsequent identification of proteinaceous effectors that are secreted by *P. brassicae* during host colonization, we developed a TurboID-based proximity labeling system in transgenic Arabidopsis Col-0, an accession of the species that is naturally susceptible to *P. brassicae* (Wang *et al*., 2023). To further refine our approach, we engineered our recombinant TurboID protein to be targeted to specific sites within host cells, namely the host nucleus, endoplasmic reticulum (ER), plasma membrane, and cytosol (Fig. 1) by incorporating well-characterized organelle-specific targeting peptides or transmembrane domains. To target recombinant TurboID (and the GFP control protein) to the nucleus, we employed the SV40 nuclear localization signal (NLS) (‘PKKKRKV’) (Kalderon *et al*., 1984), whereas a nuclear export signal (NES) from the HIV Rev protein (‘LPPLERLTL’) (Fischer *et al*., 1995) was employed to drive nuclear export, resulting in cytoplasmic accumulation of the fusion proteins. To localize our recombinant proteins to the ER, we employed the canonical C-terminal ER-retrieval motif (KDEL) (Denecke *et al*., 1991). To anchor TurboID at the inner leaflet of the plasma membrane, we considered several strategies, including fusion to the *Arabidopsis* RARE COLD INDUCIBLE2A (AtRCI2A) protein, previously used as a plasma membrane marker (Thompson & Wolniak, 2008). However, *in silico* topology predictions yielded conflicting predictions regarding whether various configurations of TurboID–RCI2A fusions would orient the labeling enzyme toward the inner (cytoplasmic-facing, intracellular position) or outer (apoplastic-facing, external position) leaflet of the membrane. Therefore, we generated two candidate TurboID fusion constructs based on domains of the well-characterized LysM receptor kinase AtCERK1 (Miya *et al*., 2007). In the first construct, the native intracellular kinase domain of CERK1 was substituted with TurboID, while retaining the upstream signal peptide, LysM motifs, and transmembrane domain (Fig. 1a), thereby positioning the labeling enzyme to face the cytoplasm. In parallel, a control reporter construct was created in which the kinase domain was replaced by GFP (Fig. 1b). The second design incorporated both a GFP reporter and the TurboID enzyme within a single recombinant protein, enabling direct visualization of subcellular localization alongside biotinylation activity. In this construct, GFP was positioned immediately downstream of the CERK1 signal peptide, replacing the LysM motifs with the fluorescent protein, while the transmembrane and juxtamembrane domains of CERK1 were retained and the kinase domain was again substituted with TurboID (Fig. 1a). For both candidate constructs, computational topology predictions indicated an intracellular orientation of the TurboID enzyme. Lastly, Flag-peptide tags (‘DYKDDDDK’) were also incorporated at the N-termini of all recombinant TurboID and GFP controls (Fig. 1a,b) to facilitate detection via Western immunoblotting, an approach that has been used to monitor TurboID accumulation previously (Qin *et al*., 2023).

**Figure 1.**
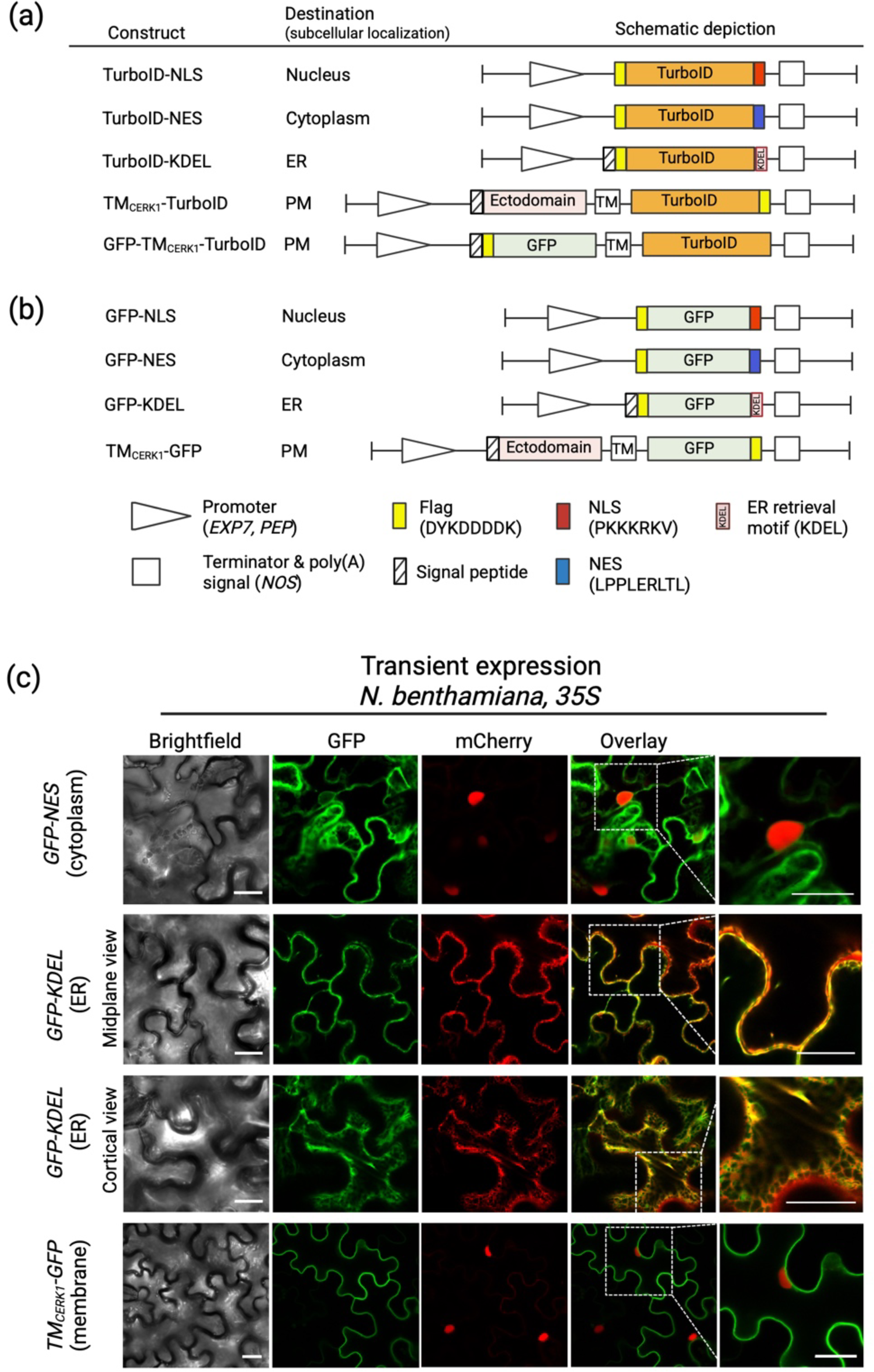
Development of a TurboID-based proximity labeling system with *in planta* subcellular specificity. (a) Schematic depiction of *TurboID-expressing* constructs and (b) *GFP*-expressing controls. (c) Transient expression of select GFP-expressing constructs in *N benthamiana* under control of *35S* promoter, co-expressed alongside subcellular markers: nuclear-localized mCherry-NLS (top and bottom panels) or ER-localized mCherry-KDEL (middle panels) Scale bars, 20um. TM, transmembrane domain. **ER,** endoplasmic reticulum. **PM,** plasma membrane.

To enable a tissue-specific resolution of *TurboID* expression, we deployed two Arabidopsis root cell type-specific promoters: an *EXPANSIN 7* promoter *(EXP7p)* that drives strong expression in trichoblast cells (Cho & Cosgrove, 2002), and a *PHOSPHOENOLPYRUVATE CARBOXYLASE* promoter (*PEPp*) that is expressed in root cortical and pericycle cells (Marquès-Bueno *et al*., 2016). These tissues represent key points of interaction between the pathogen and host during primary and secondary stages of infection, respectively (Liu *et al*., 2020), allowing us to target effectors that are specific to each of these infection stages.

To assess the accuracy of our subcellular targeting and promoter specificity prior to generating effector-fusion lines, GFP-tagged constructs were first transiently expressed in *Nicotiana benthamiana* leaves under control of the constitutive CaMV *35S* promoter and examined by confocal microscopy (Fig. 1c). Each construct that was tested exhibited the anticipated localization pattern: the NES-containing fusion localized to the cytoplasmic compartment between the vacuole and plasma membrane and signal was excluded from nuclei, consistent with active nuclear export; the ER-retained GFP colocalized with an mCherry–KDEL marker, displaying the characteristic reticulate network of the endoplasmic reticulum; and the plasma membrane–localized TM_CERK1_–GFP showed a sharp, continuous fluorescence outlining cell margins, with signal not detected in the cytoplasm or nuclei. These observations confirmed that the localization tags reliably directed proteins to their intended subcellular compartments and were suitable for downstream applications. To further evaluate promoter-driven cell-type specificity, we next generated stable Arabidopsis lines expressing the recombinant GFP fusions under control of root-specific promoters *EXP7p* and *PEPp*. Confocal imaging of roots from the corresponding transgenic lines revealed that each GFP fusion maintained the expected subcellular localization patterns (Fig. 2). Moreover, expression driven by *EXP7p* was confined to root hair and epidermal cells, whereas *PEPp* directed expression more broadly within internal cortical tissues, consistent with reported promoter activity (Marquès-Bueno et al., 2016). Collectively, these analyses validated the fidelity of our targeting strategy and confirmed that both promoter and localization elements functioned as intended, supporting the robustness of this system for downstream effector-targeting assays.

**Figure 2.**
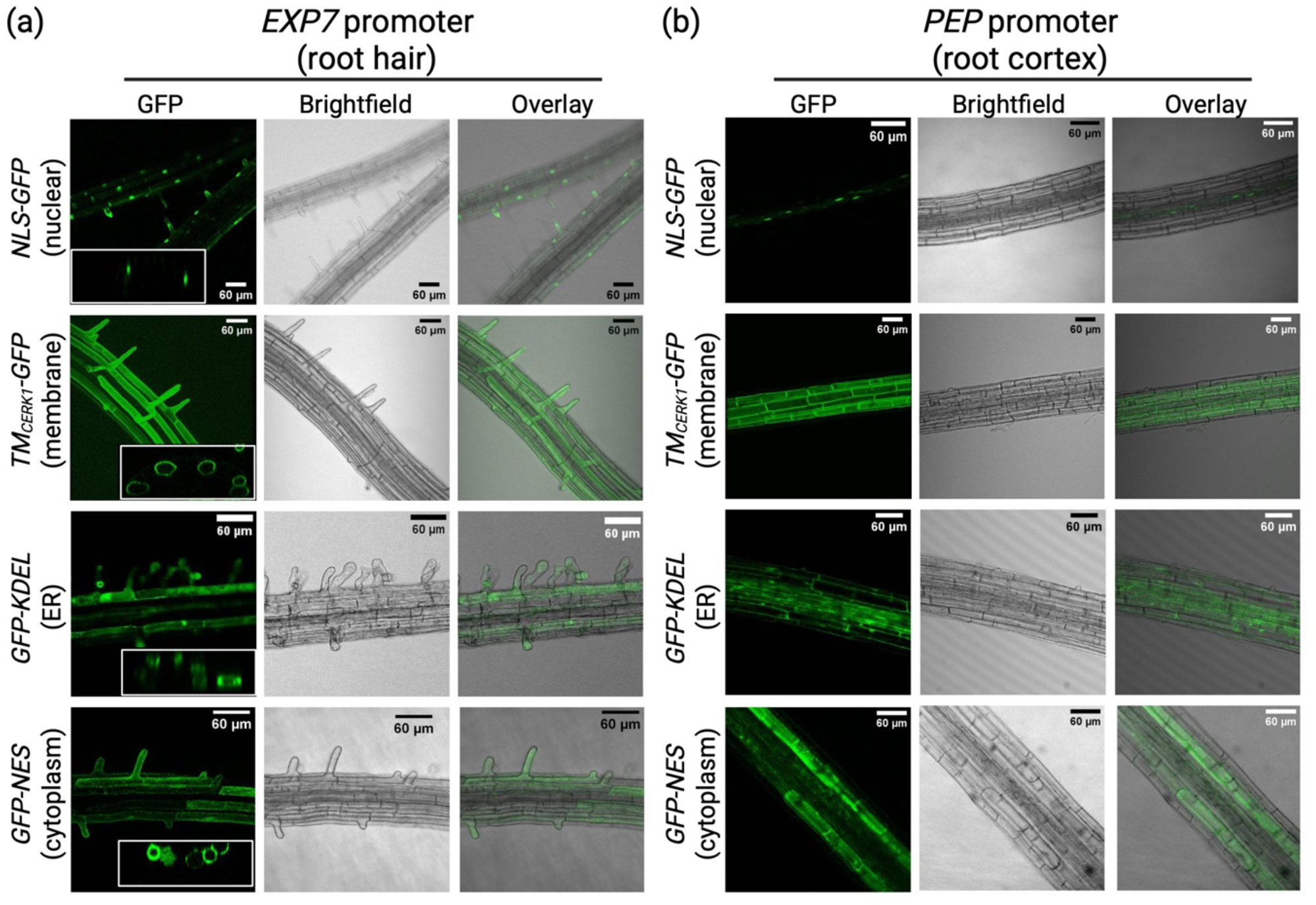
Expression of a TurboID-based proximity labeling reporters in transgenic Arabidopsis driven by root cell-specific promoters. (a) Shown is localization of GFP reporter constructs expressed under the control of (a) the *EXP7* promoter (trichoblast cells) and (b) the *PEP* promoter (cortical and pericycle cells). Scale bars, 60um.

### Development and Validation of TurboID Lines to Profile the P. brassicae Secretome

Our next objective was to identify TurboID-expressing lines exhibiting biotinylation activity suitable for downstream assays in *P. brassicae*–inoculated plants. Whereas Mair *et al*. (2019) achieved effective labeling by submerging excised seedling roots in biotin solution, our preliminary trials indicated that root tissues in intact plants could be labeled simply by watering with a biotin solution. To determine whether this non-invasive application method was effective and to establish an appropriate interval between biotin treatment and sample collection, we conducted a time-course analysis using one of the initial lines examined, *PEPp:TurboID–NLS*. Plants were watered with 20mL of 200 µM biotin, and roots were harvested for crude protein extraction at time points ranging from 0 min to 12 h post-treatment (Fig. S1). Enhanced biotinylation was detectable within 2 h and increased progressively, reaching a maximum at 8 h. Given the strong and persistent labeling observed through 12 h and the large number of plants required for subsequent experiments (∼75 per treatment group), we adopted a 10-hour biotin treatment as the standard window for most assays described hereafter, which permitted evening treatments and next-day sample processing. For each genetic construct, we successfully recovered T3 homozygous transgenic lines exhibiting clear evidence of biotinylation activity that was absent in the corresponding *GFP*-expressing control transgenic lines (Fig. 3a, b). Moreover, distinct biotinylation banding patterns were observed when TurboID was targeted to different subcellular compartments, consistent with the capture of compartment-specific proteomes. Although both membrane-localized TurboID constructs displayed labeling activity, TM_CERK1_–TurboID consistently showed higher levels of activity than GFP–TM_CERK1_–TurboID (Fig. S2). Consequently, we selected T3 homozygous lines expressing TM_CERK1_–TurboID for subsequent work.

**Figure 3.**
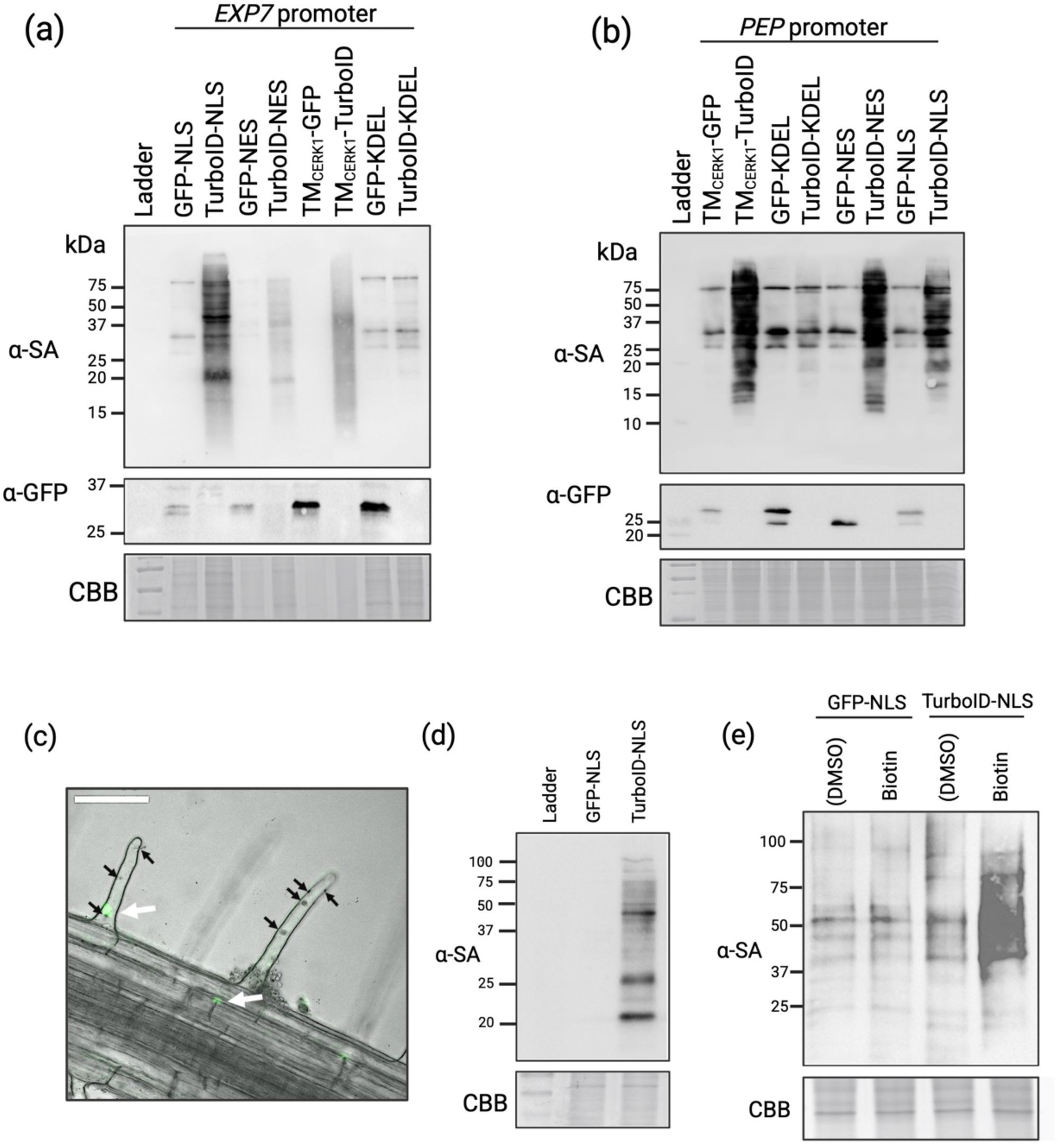
Validation of a TurboID-based proximity labeling system in transgenic Arabidopsis. Shown are biotinylation profiles of TurbolD-expressing lines and GFP-expressing controls driven by (a) the *EXP7* promoter and (b) the *PEP* promoter. (c) Confocal microscopy image showing the localization pattern of *EXP7p:GFP-NLS* in transgenic Arabidopsis roots 24 h post-inoculation with primary *P. brassicae* zoospores. White arrows indicate nuclear-localized GFP signal, and black arrows indicate encysted zoospores. (d) Biotinylation profiles of transgenic Arabidopsis lines expressing the indicated constructs under control of the EXP7 promoter, 48 hours post-inoculation with *P. brassicae* zoospores. (e) Biotinylation profiles of transgenic Arabidopsis lines expressing the indicated constructs under control of the PEP promoter, 4 weeks post-inoculation. Western blot analysis was performed on crude protein extracts from root gall tissue of plants treated with 200 µM biotin solution or a DMSO control. CBB, Coomassie Brilliant Blue stain. SA, streptavidin. Scale bar, 60um.

Our next step was to evaluate the performance of the TurboID labeling platform during primary and secondary stages of clubroot disease. To assess whether epidermal cell layer specificity of the *EXP7* promoter was maintained during primary infection, we selected the *EXP7p:GFP-NLS* line, as this construct provided bright and easily visualized fluorescence, owing to nuclear localization and concentration of the GFP signal. Confocal microscopy indicated that *EXP7p:GFP-NLS* expression remains localized to the epidermal cell layer 24- and 48-hours post-inoculation (hpi) with *P. brassicae* (Fig. 3c). We also confirmed presence of robust labeling activity in *EXP7p:TurboID-NLS* roots collected 48 hpi (Fig. 3d), and in root gall tissue harvested 4-weeks post-inoculation with *P. brassicae* (Fig. 3e). Although the structural complexity of root gall tissue prevented direct microscopic assessment of *PEP* promoter activity, previously published RNA sequencing datasets (Irani *et al*., 2018; Pérez-López *et al*., 2020) indicate that *PEP* expression levels are not detectably altered during disease progression. Collectively, these data support the use of these promoter-tag combinations for a spatially resolved and tissue-specific biotinylation of the host-pathogen proteome during *P. brassicae* infection.

### Identification of P. brassicae proteins captured within host cells during primary and secondary stages of infection

We next conducted biotinylation and pull-down assays in *P. brassicae*–infected plants, including *GFP*-expressing lines as non-TurboID expressing controls. For assays of lines expressing *PEPp*-driven transgenes (corresponding to secondary stage of disease), gall tissue was harvested from biotin-treated plants 4 weeks post-inoculation (wpi) with 10⁶ resting spores, an inoculum that consistently produces strong disease symptoms at this time point (Fig. S3). To capture the *P. brassicae* secretome during primary stage of disease, *EXP7p*-expressing roots were collected 48 hours post-inoculation (hpi) with the same spore density. Biotin-labeled proteins were isolated from crude cellular lysates by streptavidin affinity chromatography, and all samples were visualized by Western blot prior to mass spectrometry analysis (Fig. 4). In all instances, we observed evidence of robust labeling activity in TurboID samples that was absent in GFP-control samples. Importantly, we also observed strong enrichment of biotinylated proteins from beads following incubation with lysates obtained from *TurboID*-expressing samples in comparison to *GFP*-expressing samples. Lastly, analysis of unbound fractions did not reveal evidence of residual biotinylated proteins, confirming the effectiveness of the affinity purification.

**Figure 4.**
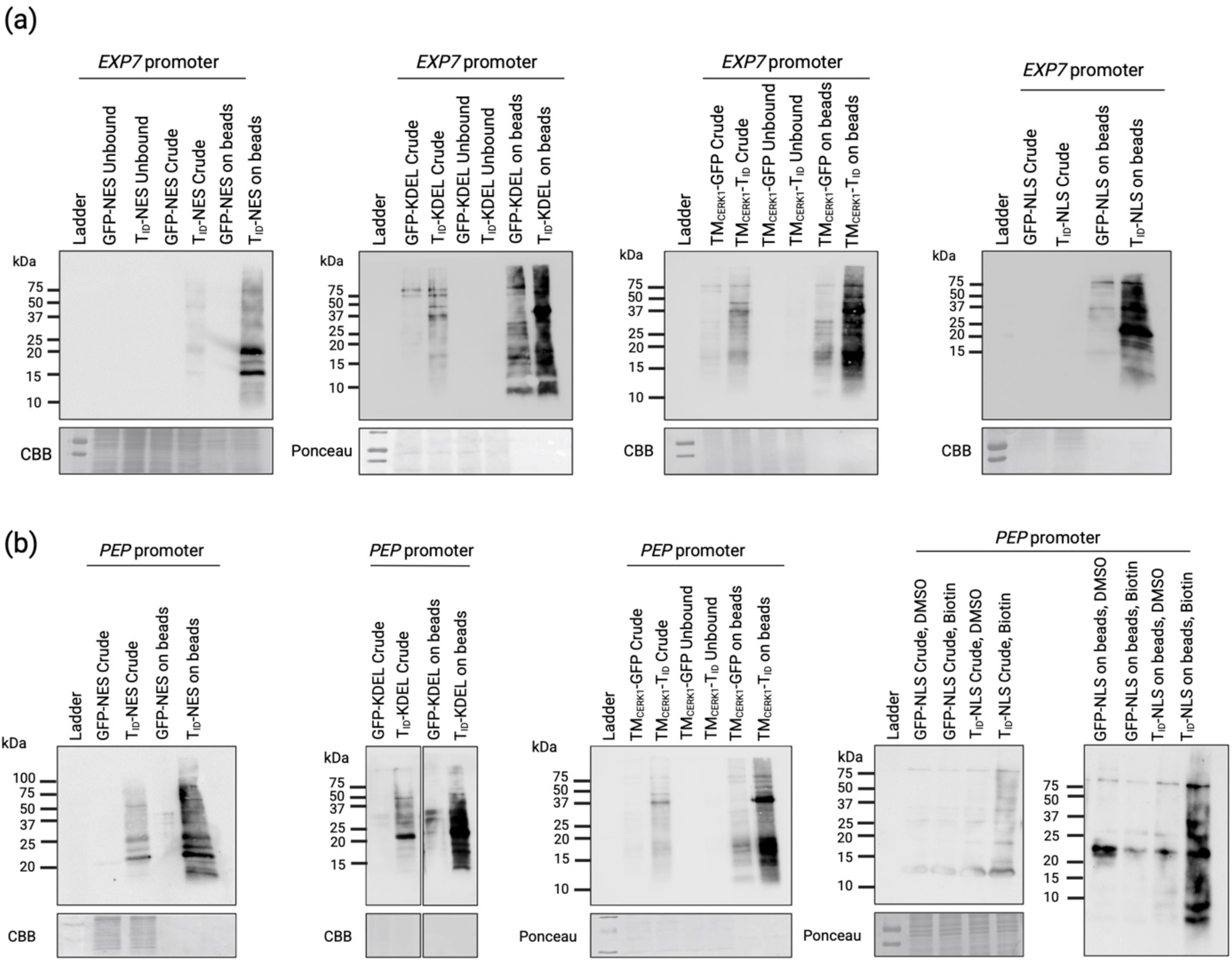
Affinity purification of biotinylated proteins recovered from transgenic *Arabidopsis* plants following infection with *P. brassicae.* Shown are biotinylation profiles of TurbolD-expressing lines and GFP-expressing controls driven by (a) the *EXP7* promoter and (b) the *PEP* promoter. Samples were collected 48 hours (a) or 4 weeks (b) post-inoculation. In all cases, blots were probed with anti-streptavidin antibody. CBB, Coomassie Brilliant Blue stain. Ponceau, Ponceau stain.

In total, we identified 106 *P. brassicae* proteins in *EXP7p:TurboID* lines and 281 in *PEPp:TurboID* lines (Table S2). Both lists were then curated to remove endogenously biotinylated host carboxylases and highly abundant non-specific interactors that have been demonstrated to frequently contaminate affinity purifications (false positives corresponding to the so-called “CRAPome”; (Mellacheruvu *et al*., 2013; Petre *et al*., 2015, 2021; Van Leene *et al*., 2015). We then defined a high-confidence candidate effector set consisting of *P. brassicae* proteins predicted to contain an N-terminal signal peptide but no transmembrane domains, and that were detected in at least one TurboID-expressing line (Table 1). Among the proteins represented in our high-confidence list was PbBSMT, a salicylic acid methyltransferase that represents one of the best-characterized *P. brassicae* effectors to date. Interestingly, PbBSMT was detected in *PEPp* samples derived from membrane- and ER-localized TurboID lines and was absent from those expressing nuclear- or cytoplasmic-localized TurboID. Also present were three *P. brassicae* proteins (CEO97005, CEO97020, and CEP01791) previously described as effector candidates PbPE8, PbPE35, and PbPE48, respectively (Hossain et al., 2021). These proteins were highly abundant in TurboID datasets corresponding to the secondary stage of clubroot disease yet were absent from those representing primary infection.

**Table 1.** High-confidence *P. brassicae effector* candidates.

| GenBank<br>Accession | Annotation (UniProt) | Sample(s) | Previous Description |
| --- | --- | --- | --- |
| CEP01733.1 | Uncharacterized protein<br>(terpenoid synthase superfamily) | EXP7-NLS | None |
| CEP03710.1 | Protein kinase domain-containing<br>protein | EXP7-NLS,<br>EXP7-NES,<br>PEP-NES | PbPE23;<br>(Hossain et al., 2024) |
| CEO97005.1 | Uncharacterized protein | PEP-NLS<br>PEP-NES,<br>PEP-CERK | PbPE8;<br>(Hossain et al., 2021) |
| CEP01791.1 | Histidine kinase/HSP90-like<br>ATPase domain-containing<br>protein | PEP-NLS<br>PEP-KDEL,<br>PEP-CERK | PbPE48;<br>(Hossain et al., 2021) |
| CEO98662.1 | Protein kinase domain-containing<br>protein | PEP-NLS | None |
| CEO97020.1 | Uncharacterized protein | PEP-NLS<br>PEP-NES,<br>PEP-KDEL,<br>PEP-CERK | PbPE35;<br>(Hossain et al., 2021) |
| CEO97827.1 | Uncharacterized protein (low<br>complexity/disordered structure) | PEP-NLS<br>PEP-NES,<br>PEP-KDEL,<br>PEP-CERK | None |
| CEO98333.1 | Uncharacterized protein | PEP-NES | None |
| CEO95736.1 | Serine/threonine-protein<br>phosphatase (EC 3.1.3.16) | PEP-KDEL | None |
| CEO94659.1 | Benzoic acid/salicylic acid<br>methyltransferase | PEP-KDEL<br>PEP-CERK | PbBSMT;<br>(Ludwig-Müller et al.,<br>2015) |
| CEO95605.1 | Uncharacterized protein<br>(predicted ATP-dependent<br>protein folding chaperone) | PEP-KDEL<br>PEP-CERK | None |
| CEP02998.1 | Band 7 domain-containing protein | PEP-CERK | None |
| CEP02396.1 | Protein kinase domain-containing protein | PEP-CERK | SSPbP22<br>(Pérez-López et al., 2020) |

We also identified PbPE23, a protein with a predicted Serine/Threonine kinase domain that had likewise been characterized as an effector candidate (Hossain et al., 2024). PbPE23-derived peptides were recovered from both cytoplasmic (TurboID-NES) and nuclear (TurboID-NLS) datasets under expression driven by both the *EXP7* and *PEP* promoters. In addition to PbPE23, only a single other secreted protein was detected during primary infection—CEP01733 (PBRA_008675)—which encodes a predicted isoprenoid-biosynthesis–related domain and has not yet been experimentally investigated.

Samples obtained from *PEPp-*expressing transgenes yielded a greater number of high confidence effector candidates (Table 1), with a total of 12 secreted proteins detected in *PEPp:TurboID-NLS* (5), *PEPp:TurboID-NES* (5), *PEPp:TurboID-KDEL* (6) and *PEPp::TM_CERK1_-TurboID* (8), with a subset of proteins that are shared between two or more datasets. The *PEPp:TurboID-NLS* dataset yielded two proteins with predicted kinase domains; PbPE48/CEP01791, identified as a predicted histidine kinase, and CEO98662, annotated as a ‘Protein kinase domain-containing protein’ due to the presence of a kinase catalytic domain at amino acids (47-279). Four candidates are annotated simply as “uncharacterized proteins” and lack detectable conserved domains. Notably, one of these (CEO97827) contains a low-complexity, intrinsically disordered region (IDR), a feature frequently associated with pathogen effectors (Marín *et al*., 2013; Chepsergon & Moleleki, 2023).

## Discussion

*P. brassicae* belongs to the class *Phytomyxea* within the supergroup *Rhizaria—*a relatively understudied lineage of eukaryotes that includes other plant parasites such as *Spongospora subterranea,* the causal agent of potato powdery scab (Schwelm *et al*., 2015). This group of microbial eukaryotes is only distantly related to the well-characterized fungal and oomycete pathogens that inform much of our current understanding of effector biology. When this project was conceived (circa 2019), it remained uncertain whether *P. brassicae* might employ noncanonical effector secretion mechanisms beyond the classical N-terminal signal peptide–dependent pathway. We therefore developed an unbiased, experimental proximity-labeling approach to complement existing *in silico* secretome predictions, reasoning that this strategy could capture both conventionally and unconventionally secreted proteins, should alternative export routes be active in *P. brassicae.* Moreover, we aimed to comprehensively profile its secretome across distinct host subcellular environments. To this end, we targeted the proximity labeling enzyme TurboID to four key host compartments - the nucleus, cytosol, plasma membrane, and ER - which are frequent destinations for effector localization in other plant–pathogen systems (Caillaud *et al*., 2012; Khan *et al*., 2018). The inclusion of ER- and membrane-localized TurboID reporters was further motivated by prior preliminary evidence that *P. brassicae* effectors target the host endomembrane system during infection (Hossain *et al*., 2021).

Since that time, several independent studies have provided compelling evidence that *P. brassicae* proteins are indeed secreted via the classical (ER)–Golgi route, and encode a canonical N-terminal signal peptide that is functional in yeast (Chen *et al*., 2019; Djavaheri *et al*., 2019; Yu *et al*., 2019; Pérez-López *et al*., 2020; Hossain *et al*., 2021; Feng *et al*., 2024; Yang *et al*., 2024). In light of this emerging consensus, we mined our proximity labeling results to extract all *P. brassicae* proteins detected in our TurboID-expressing samples that also encode a predicted signal peptide (Table 1). This filtering step substantially narrowed our list of 339 candidate genes (Table S1) to a much smaller high-confidence subset, representing those most likely to function as bona fide secreted effectors.

Included among our high-confidence proteins were several previously investigated effector candidates, most notably PbBSMT, one of the best-characterized *P. brassicae* effectors. PbBSMT is a SABATH-type methyltransferase that catalyzes the conversion of SA to methyl salicylate (MeSA), thereby reducing the accumulation of bioactive SA and suppressing host defense signaling (Ludwig-Müller *et al*., 2015; Bulman *et al*., 2019; Djavaheri *et al*., 2019). Consistent with earlier data showing induction of *PbBSMT* expression during secondary infection (Ludwig-Müller *et al*., 2015; Rolfe *et al*., 2016), this protein was detected exclusively in *PEPp*-expressing lines, implying a function that is primarily associated with the secondary phase of disease. Strikingly, PbBSMT was only recovered from *PEPp:TurboID-KDEL* and *PEPp:TMCERK1-TurboID* datasets, suggesting this effector accumulates in close proximity to the host ER and plasma membrane. Transmission electron microscopy has shown that *P. brassicae* secondary zoospores and plasmodia often reside adjacent to, and even traverse, plasmodesmata connecting neighbouring cortical cells (Djavaheri *et al*., 2023). Moreover, there exists evidence that at least one other *P. brassicae* effector (PbPE13) localizes to plasmodesma (Hossain *et al*., 2021), where it may play a role in suppressing host immunity at this location, further supporting a model in which plasmodesmata represent key sites of plasmodial interaction and activity during infection. As SA regulates plasmodesmatal permeability through callose deposition and closure (Wang *et al*., 2013), a locally acting SA-methylating enzyme could plausibly function to actively influence intercellular connectivity. One interpretation of our data, in other words, is a model whereby PbBSMT is secreted to the vicinity of plasmodesmatal and host:pathogen interface membranes, where the enzyme acts to impair SA-mediated signalling at these sites. This interpretation is supported by the known enzymatic function of PbBSMT (Ludwig-Müller *et al*., 2015), its enrichment in membrane- and ER-associated TurboID datasets, and FISH analyses showing *PbBSMT* transcript accumulation at this host–pathogen interface (Badstöber *et al*., 2020). Collectively, these observations suggest that *P. brassicae* secretes PbBSMT in a spatially targeted manner to enhance intracellular continuity across infected cortical tissues, while locally suppressing salicylic acid–dependent defense responses to facilitate pathogen spread.

We likewise recovered the previously characterized kinase effector protein PbPE23 in our datasets, which has been shown to induce necrosis when expressed in both host and non-host plants (Hossain et al., 2024). Hossain and colleagues reported a nucleocytoplasmic localization for this protein *in planta*, consistent with our finding that PbPE23 was detected exclusively in TurboID-NES and TurboID-NLS samples and was absent from TurboID-KDEL and TM_CERK1_-TurboID datasets. Notably, we detected PbPE23 in both *EXP7p*- and *PEPp-*driven transgenic lines, suggesting that its activity spans both primary and secondary stages of infection. Although Hossain et al. observed increased PbPE23 transcript accumulation during the transition from primary to secondary infection, our data indicate that the corresponding protein may be present earlier in the disease cycle than previously reported.

In addition to PbPE23, we only recovered one other secreted *P. brassicae* protein in our *EXP7p* datasets (CEP01733), suggesting that either *P. brassicae* effectors play a less active role during the primary stage of infection, or that our experimental set up may not have been conducive to the capture of such proteins. One possibility to account for the low number of secreted proteins in *EXP7p* datasets is that effector deployment during the primary root-hair infection stage may be limited or highly transient. This observation could reflect biological differences in the infection process, as primary infection is largely confined to individual root hairs and may rely more on mechanical or physiological compatibility than on extensive effector-mediated suppression of immunity. Alternatively, the result may indicate technical constraints of our experimental system that limited the recovery of secreted proteins at this stage, particularly given that whole root systems were harvested rather than isolated root hairs — a necessary compromise due to the impracticality of obtaining sufficient quantities of root-hair material for affinity purification. This may have reduced sensitivity and diluted signals specific to the primary infection sites.

Among the proteins detected during primary infection, CEP01733 (PBRA_008675) is predicted to be a secreted protein with a conserved isoprenoid/terpenoid biosynthesis domain. Isoprenoids form a major class of metabolites involved in both primary and defense-related pathways, including precursors of phytohormones such as gibberellins, abscisic acid, brassinosteroids and cytokinins (Tarkowská & Strnad, 2018). Manipulation of host isoprenoid metabolism is a recurrent feature of gall-forming and other biotrophic pathogens, which frequently reprogram hormone homeostasis to promote cell hypertrophy and dampen immune responses (Vañó *et al*., 2023). Transcriptomic data further indicate that *P. brassicae* colonization of root hair cells at this time point coincides with elevated expression of host genes involved in terpenoid biosynthesis (Zhao *et al*., 2017), suggesting an early perturbation of this metabolic network. A secreted *P. brassicae* enzyme possessing terpenoid biosynthetic activity could plausibly influence this process, for example, by locally modifying metabolite availability at the host–pathogen interface. Taken together, the detection of a secreted *P. brassicae* protein containing a conserved isoprenoid/terpenoid biosynthesis domain during primary infection, alongside evidence for host terpenoid pathway activation at the same stage, supports a model in which enhanced terpenoid metabolism contributes to establishing early compatibility between parasite and host.

Recovery of multiple *P. brassicae* secreted proteins containing predicted kinase domains and one protein predicted to be a protein phosphatase raises the intriguing possibility that this pathogen encodes effectors capable of directly targeting host signalling processes via phosphorylation. Phosphorylation is critical in a multitude of signaling pathways that are crucial to host cells that may be beneficial for Plasmodiophora to manipulate, for example, cell cycle initiation or host defence signalling. Kinase-like effectors have been described in bacterial and oomycete phytopathogens, where they target immunity-associated signalling pathways (Teper *et al*., 2018; Ai *et al*., 2021). The detection of such proteins in our proximity-labeling experiments suggests that *P. brassicae* may employ a similar strategy to impair host defence, potentially through phosphorylation-mediated manipulation of plant signaling pathways. While the kinase activity of PbPE23 has been experimentally verified (Hossain *et al*., 2024), the kinase domain in CEO98662 (PBRA_006776) appears to lack canonical catalytic motifs of common secreted enzyme classes (e.g., HRD, DFG, VAIK motifs of Ser/Thr/Tyr kinases), suggesting a non-enzymatic effector or a divergent enzyme whose function will require further experimental validation.

In conclusion, this study provides new insights and establishes a foundation for advancing our understanding of how *P. brassicae* - a representative of the poorly characterized *Rhizaria* - infects and causes disease in its plant hosts. Our proximity labeling approach provides the first experimental view of the *P. brassicae* secretome within the host cellular environment, yielding a high-confidence dataset that includes both established and previously uncharacterized effectors. This dataset represents a valuable community resource for guiding future research endeavours into effector function and host target identification. More broadly, this work demonstrates that proximity labeling can overcome long-standing barriers associated with studying obligate biotrophs, providing a scalable strategy for experimentally defining the effector repertoires of other intracellular plant pathogens.

## Acknowledgements

Funding for this work was provided to Drs. Christopher Todd and Allyson MacLean by the Canola Council of Canada and the Western Grains Research Foundation (CARP2020.07), and to Dr. Kris Kalinger through a Natural Sciences and Engineering Research Council of Canada (NSERC) Postdoctoral Fellowship.

## Author contributions

AMM and CDT conceived the study and secured funding. EM performed cloning and generated transgenic Arabidopsis lines. MN conducted the initial screening of transgenic lines for TurboID ligase activity, assessed biotinylation in root galls, and carried out time-course experiments and initial optimization of inoculation protocols. MMH performed an initial microscopic localization assessment in Arabidopsis and subcellular localization assays *Nicotiana benthamiana*. Confocal imaging of transgenic Arabidopsis roots was conducted by EGB, using materials provided by KK. KK performed all affinity purification experiments, post-acquisition mass spectrometry data analyses, and identified the best-performing transgenic lines for secretome analysis. KK also prepared all blots and gels shown in the figures, except those from the time-course and root gall experiments, and generated draft figures. Mass spectrometry analyses were conducted by MT and RGU. AMM wrote the first draft of the manuscript and assembled the accompanying figures. All authors contributed to manuscript revision and approved the final version.

## Data availability

Data will be made available upon request.

Mass spectrometry data have been deposited to the ProteomeExchange Consortium (http://proteomecentral.proteomexchange.org) via the PRoteomics IDEntification Database (PRIDE; https://www.ebi.ac.uk/pride/) partner repository with the data set identifier PXD069805.

## Supporting Information

Table S1. List of oligonucleotide primers used to create proximity labeling and reporter constructs

Table S2. Total *P. brassicae* proteins detected in *EXP7* and *PEP* TurboID datasets

**Figure S1.**
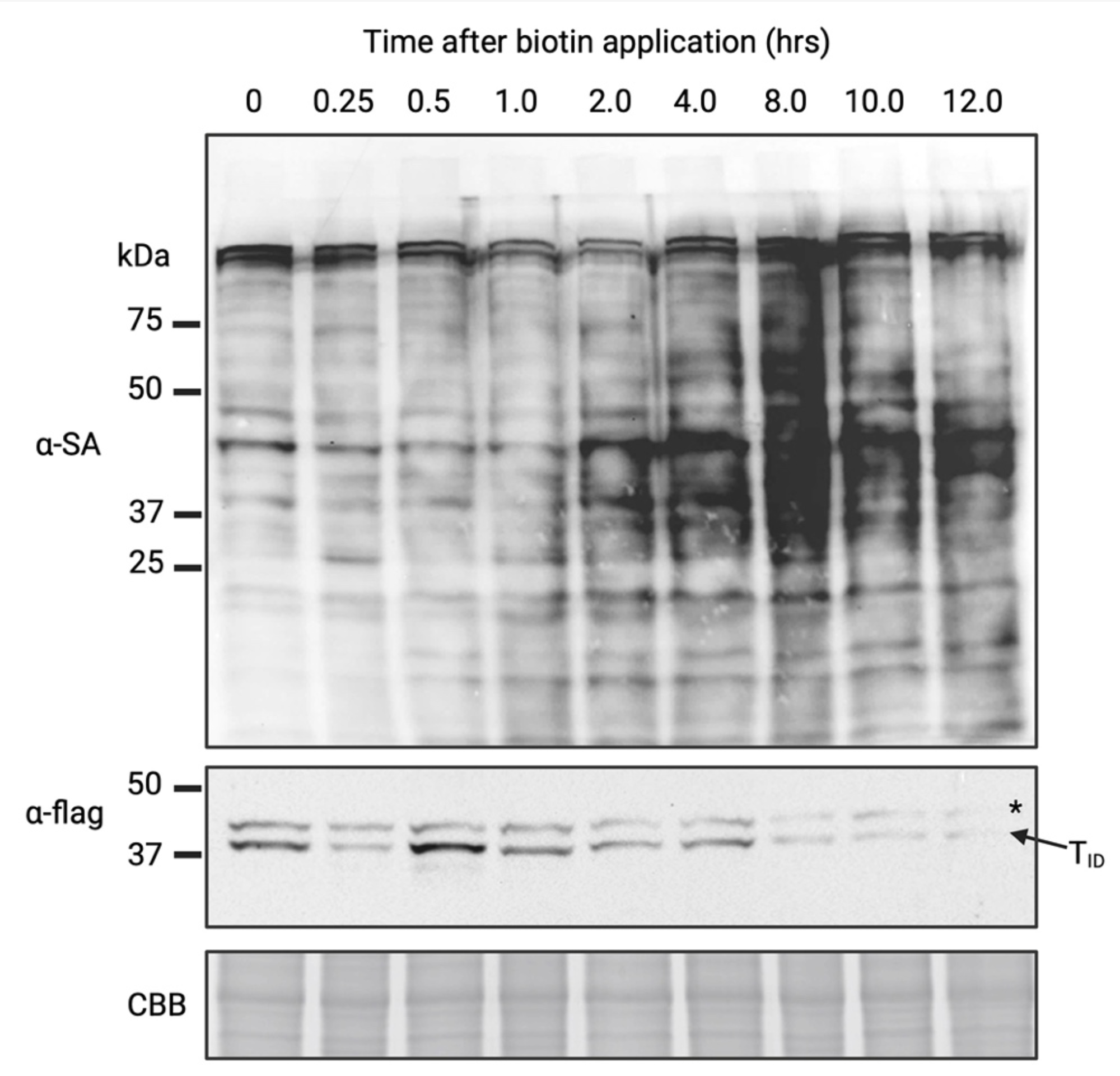
Time course of total protein biotinylation in transgenic Arabidopsis roots following exogenous biotin treatment. Arabidopsis Col-0 plants expressing FLAG-tagged TurbolD:NLS expressed under control of the *PEP* promoter were treated by watering with 200 µM biotin solution, with root systems collected at the indicated time points (0-12 h). **Upper panel:** Western blot probed with anti-streptavidin-HRP to detect biotinylated proteins. **Middle panel:** Anti-FLAG immunoblot to assess TurbolD abundance. **Lower panel:** Coomassie-stained gel showing total protein as a loading control. The asterisk denotes a non-specific band commonly observed in anti-FLAG blots of Arabidopsis protein extracts.

**Figure S2.**
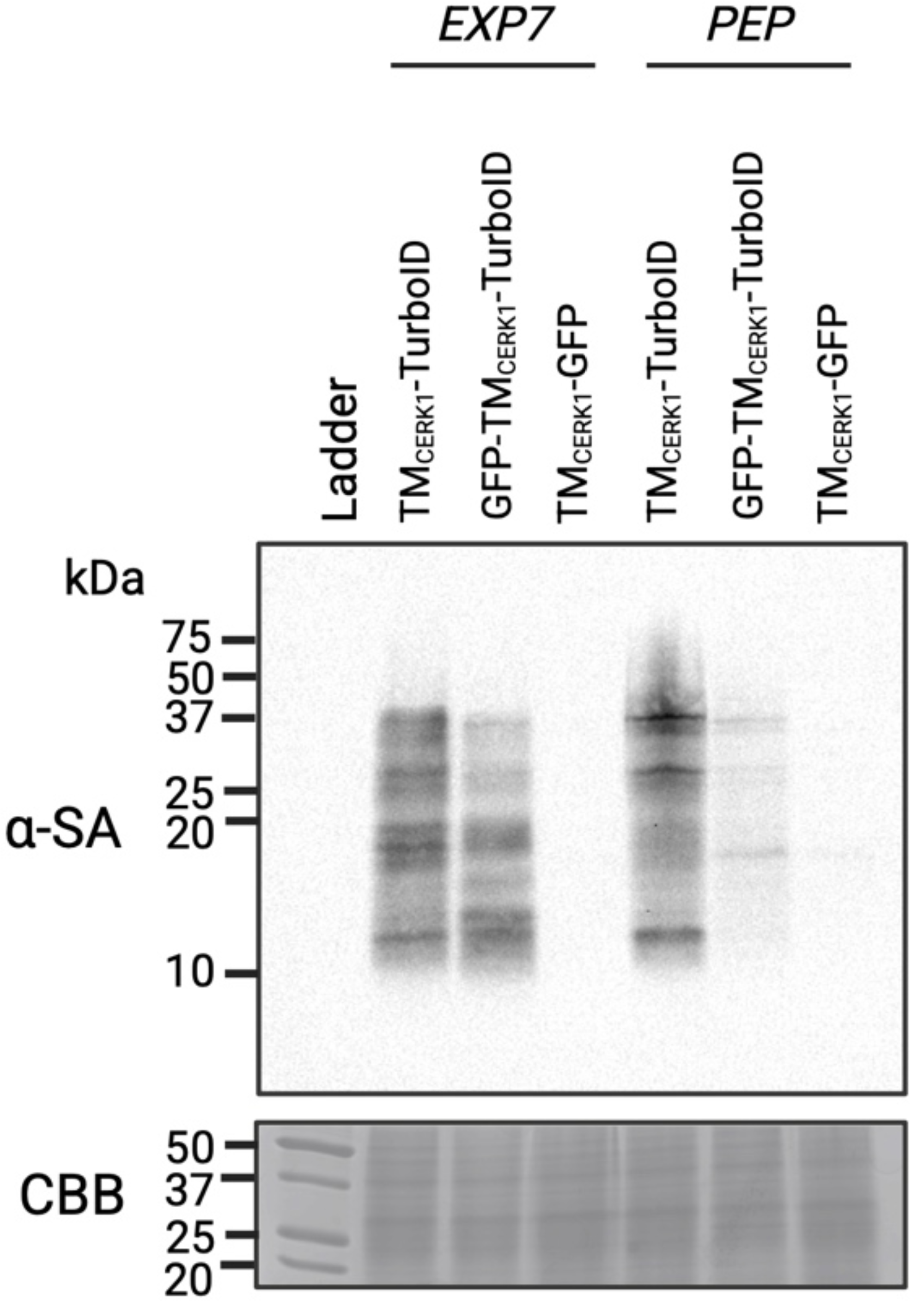
Comparison of membrane-localized TurboID constructs for biotin ligase activity. Transgenic *Arabidopsis* lines expressing the indicated constructs were treated with 200 µM biotin solution and incubated for 10 h prior to sample collection. Plants were grown in the absence of *Plasmodiophora brassicae* for two weeks prior to biotin treatment.

**Figure S3.**
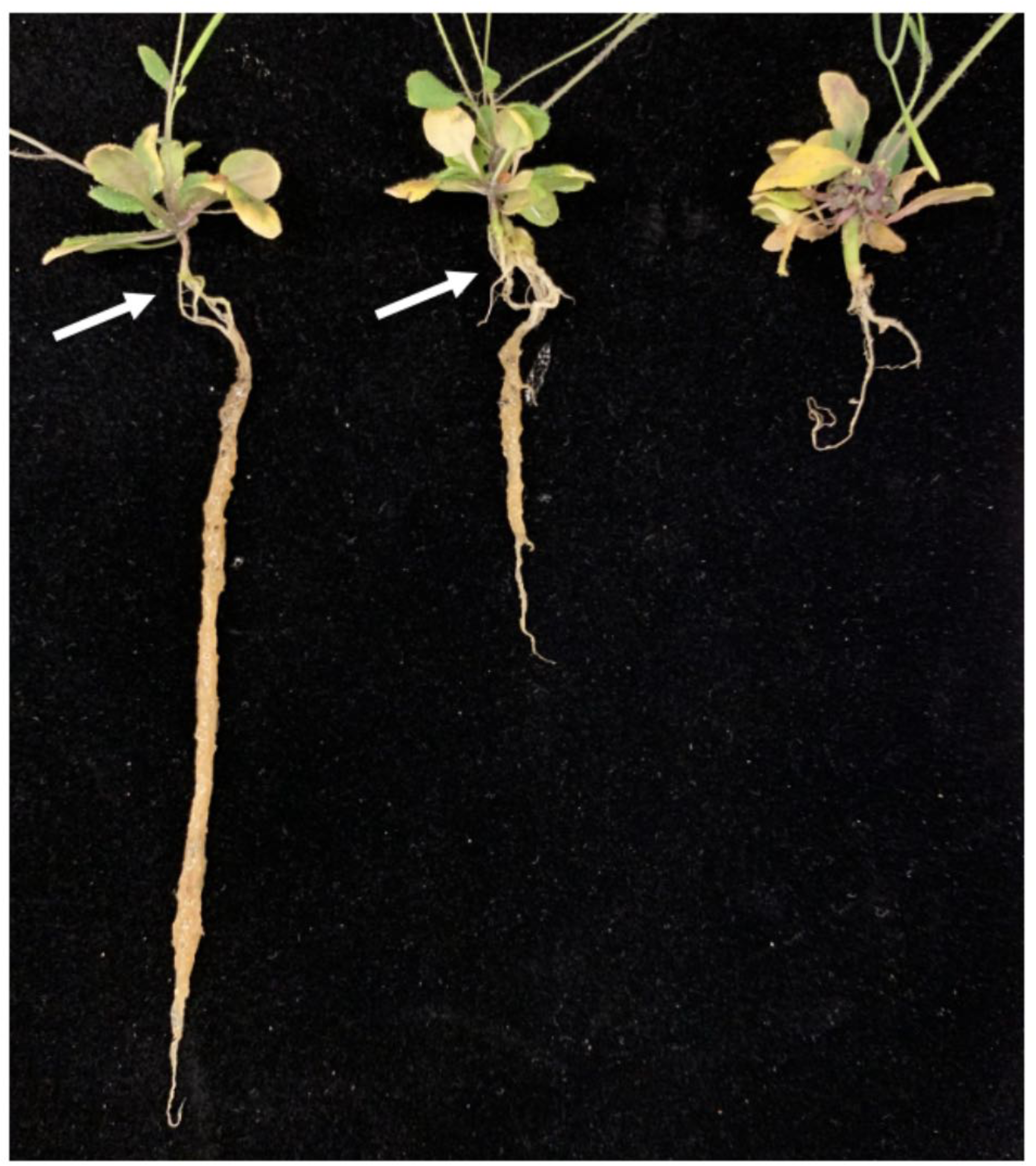
Infection progress at 4 weeks post-inoculation (wpi) in *Arabidopsis thaliana* Col-0 plants inoculated with varying concentrations of *Plasmodiophora brassicae* resting spores. Plants were inoculated 12 days after germination. Inoculation with 10^4^ resting spores resulted in mild disease symptoms, including leaf chlorosis, wilting, and small galls near the hypocotyl (white arrow), while the remainder of the root system remained intact. Plants inoculated with 10^6^ spores developed larger galls and exhibited pronounced stunting of root growth. At the highest inoculum (10^8^ spores), by 4 wpi only thickened, diseased roots remained, with extensive chlorosis, wilting, and destruction of the root system.

