## Supplemental Table S1 for "TurboID-based proximity labeling enables *in vivo* mapping of *Plasmodiophora brassicae* secretome in Arabidopsis"

Table S1. List of Oligonucleotide Primers Used to Create Proximity Labeling and Reporter Constructs

| **Primer Name** | **Sequence (5′ → 3′)** | **Purpose / Notes** |
| --- | --- | --- |
| EM-F1 | ATGGACTACAAGGACGACGATGACAAAGTGAGCAAGGGCGAGGAG | Adds FLAG epitope (eGFP fragment) |
| EM-R1 | TCACTACAGGGTCAGGCGCTCCAGGGGAGGCAGCTTGTACAGCTCGTCCATG | Adds NES epitope + 2 stop codons |
| EM-F2 | ATGGACTACAAGGACGACGATGACAAAGCTAGCAAAGACAATACTGTG | Amplifies TurboID-NES; adds FLAG epitope |
| EM-R2 | TCACTACAGGGTCAGGCGCTCCAG | Reverse for TurboID-NES fragment |
| EM-F3 | CGGTACCCGGGGATCCTTAAAAGCGTTTTTTTTTATTCTGAG | EXP7 promoter (adds vector overlap) |
| EM-R3 | GTCGTCCTTGTAGTCCATTCTAGCCTCTTTTTCTTTATTCTT | Overlap with FLAG sequence |
| EM-F4 | ATGGACTACAAGGACGACG | Reamplifies FLAG-tagged fragments; vector overlap |
| EM-R4 | GGTCACCTGTAATTCACACTCACTACAGGGTCAGGCG | Vector (NOST region) overlap |
| F13 | CGGGGATCCTCTAGATAAAAGCGTTTTTTTTTATTCTGAG | EXP7 promoter cloning (XbaI site) |
| R13 | CTGTAATTCACACGTGTCTAGCCTCTTTTTCTTTATTCTT | EXP7 promoter cloning (PmlI site) |
| F14 | CGGGGATCCTCTAGAATCAAGACCATCTGTAATCT | PEP promoter cloning (XbaI site) |
| R14 | CTGTAATTCACACGTGGGTTTTGGCTAATGTGATTG | PEP promoter cloning (PmlI site) |
| EM-F8 | CACATTAGCCAAAACCCACATGGACTACAAGGACGACG | Adds overlap with PEP promoter |
| EM-R8 | TCACTAGAGTTCATCCTTCTTGTACAGCTCGTCCATG | Adds KDEL-2 stops |
| EM-F10 | AGAAAAAGAGGCTAGACACATGGACTACAAGGACGACG | Adds overlap for EXP7 promoter |
| EM-R10 | GTCACCTGTAATTCACACTCACTAGAGTTCATCCTTCT | Adds overlap with vector (NOST) |
| EM-R9a | TCACTAGAGTTCATCCTTCTGCAGCTTTTCGGCAGA | Adds KDEL-2 stops to TurboID |
| EM-F11 | ACATTAGCCAAAACCCACATGGACTACAAGGACGACG | Adds overlap for PEP promoter |
| EM-F46 | TCCACCATGAAGCTAAAGATTTCTCTAA | Amplifies CERK1 CDS; adds Kozak sequence |
| EM-R46 | CACGGTAATGGTTCTCGTCA | CERK1 reverse primer |
| EM-F49 | AGAAAAAGAGGCTAGACACTCCACCATGAAGCTAAAGAT | Overlap with EXP7 promoter; CERK1 fragment |
| EM-R49 | ATTGTCTTTCAGCTCTGCATAGTAAACAG | Overlap with TurboID fragment |
| EM-F50 | TGCAGAGCTGAAAGACAATACTGTGCCTCT | Overlap with CERK1 fragment |
| EM-R50 | CTACTATTTGTCATCGTCGTCCTTGTAGTCCTTTTCGGCAGACCGCAG | Adds FLAG + 2 stops |
| EM-R51 | GTCACCTGTAATTCACACCTACTATTTGTCATCGTCGT | Adds vector overlap (NOS-T region) |
| EM-R52 | CTTGCTCACTTTGTCATCGTCGTCCTTGTAGTCCATGCACTTAGATTCCACGGC | Adds FLAG + overlap with GFP |
| EM-F53 | CGATGACAAAGTGAGCAAGGGCGAGGA | Overlap with FLAG epitope (GFP fragment) |
| EM-R53 | ACCAACACCCTTGTACAGCTCGTCCATG | Overlap with CERK1 fragment |
| EM-F54 | CTGTACAAGGGTGTTGGTGCTGGAGTT | Overlap with GFP fragment |
| EM-R54 | ATTGTCTTTCAGCTCTGCATAGTAAACAG | Overlap with TurboID |
| EM-F55 | GCAGAGCTGAAAGACAATACTGTGCCTCT | Overlap with CERK1 |
| EM-R55 | GTCACCTGTAATTCACACCTACTACTTTTCGGCAGACCGCAG | Overlap with vector (NOS-T region) |
| EM-F57 | CAGAGCTGGTGAGCAAGGGCGAGGA | Overlap with CERK1 fragment |
| EM-R57 | CTACTATTTGTCATCGTCGTCCTTGTAGTCCTTGTACAGCTCGTCCATG | Adds FLAG + 2 stops codons |
| EM-F58 | ACATTAGCCAAAACCCACTCCACCATGAAGCTAAAGAT | Adds overlap with PEP promoter |
| EM-R56 | TTGCTCACCAGCTCTGCATAGTAAACAG | Overlap with GFP fragment |
| EM-F10 | AGAAAAAGAGGCTAGACACATGGACTACAAGGACGACG | Adds overlap for EXP7 promoter |
| EM-R10 | GTCACCTGTAATTCACACTCACTAGAGTTCATCCTTCT | Adds overlap with vector (NOST) |
| EM-R9a | TCACTAGAGTTCATCCTTCTGCAGCTTTTCGGCAGA | Adds KDEL-2 stops to TurboID |
| EM-F11 | ACATTAGCCAAAACCCACATGGACTACAAGGACGACG | Adds overlap for PEP promoter |
| EM-F46 | TCCACCATGAAGCTAAAGATTTCTCTAA | Amplifies CERK1 CDS; adds Kozak sequence |
| EM-R46 | CACGGTAATGGTTCTCGTCA | CERK1 reverse primer |
| EM-F49 | AGAAAAAGAGGCTAGACACTCCACCATGAAGCTAAAGAT | Overlap with EXP7 promoter; CERK1 fragment |
| EM-R49 | ATTGTCTTTCAGCTCTGCATAGTAAACAG | Overlap with TurboID fragment |
| EM-F50 | TGCAGAGCTGAAAGACAATACTGTGCCTCT | Overlap with CERK1 fragment |
